# Spatial Navigation Training Enhances Large-Scale and Small-Scale Spatial Abilities through Different Neural Mechanisms

**DOI:** 10.64898/2025.12.18.695072

**Authors:** Jin Yu, Mengxia Yu, Yiying Song, Siyuan Hu

## Abstract

Elucidating the relationship between large-scale and small-scale spatial abilities is fundamental to advancing our understanding of spatial cognition, with training transfer effects across tasks offering a direct means of exploration. This study investigated how large-scale spatial navigation training influences both large- and small-scale spatial abilities and their underlying neural mechanisms. Participants completed 20 days of real-world campus navigation training and performed large-scale (distance judgment) and small-scale (paper folding) spatial tasks before and after training. The training group showed significant improvements in both tasks, whereas the control group did not. Brain imaging revealed increased activation in the right middle frontal gyrus (MFG) and bilateral posterior cingulate cortex (PCC) during large-scale task performance after training. In contrast, improvements in the small-scale task were associated with reduced deactivation in the right postcentral gyrus (PoCG), right precuneus, and left superior temporal gyrus (STG). Overall, these findings indicate that large-scale navigation training enhances spatial ability across scales through distinct neural mechanisms, supporting the partial dissociation model and highlighting the contribution of negatively activated regions to spatial processing.

## 1. Introduction

Spatial abilities, a crucial nonverbal component of intelligence, can be divided into large-scale and small-scale types (Hegarty et al., 2006). Large-scale spatial abilities are reflected in tasks involving broad environments, such as spatial navigation (Wolbers & Wiener, 2014) and route planning (Farr et al., 2012). These tasks often require observers to use their own position as a reference to navigate the environment (Evans, 1980). In contrast, small-scale spatial tasks typically occur in controlled small-scale settings, such as mental rotation (Wexler et al., 1998), paper folding (Shepard & Feng, 1972), and embedded figure tests (Walter & Dassonville, 2011). These tasks require the observer to visualize objects from a fixed perspective and mentally manipulate the relationships between different parts of the objects (Kozhevnikov & Hegarty, 2001). Together, these two spatial ability types play essential but distinct roles in how we perceive and interact with the environment.

A key unresolved issue is the relationship between large- and small-scale spatial abilities. Hegarty et al. (2006) proposed four possible models: they may be completely unrelated (total dissociation model); partially related (partial dissociation model); represented by a single, unified ability (unitary model); or jointly influenced by a third variable (mediation model). Among these, the partial dissociation model has received the most empirical support, with some studies reporting correlations ranging from 0.50 to 0.87 (Kozhevnikov & Hegarty, 2001; Hegarty et al., 2006; Likhanov et al., 2022). Other studies, however, have reported minimal shared variance between the two abilities (Wang et al., 2014; Quaiser-Pohl et al., 2004). Overall, these mixed findings suggest that some degree of association exists between large- and small-scale spatial abilities.

Further evidence for this association comes from spatial training transfer—enhancing one may benefit the other. Previous research has shown that training effects can extend beyond the practiced task (Uttal et al., 2013). Some studies have examined whether such transfer extends across spatial scales. For instance, Jansen et al. (2010) trained participants on a manual rotation task and reported that the training group made smaller direction estimate errors in a virtual environment, indicating that small-scale training transferred to large-scale spatial performance. This transfer may occur because the two types of tasks share some common cognitive components. With respect to large-scale spatial training, some studies have shown that navigation training improves task performance as well as associated neural activation (Auger et al., 2017; Wenger et al., 2012). Beyond effects within large-scale contexts, training on large-scale skills such as map reading may also improve small-scale mental rotation (Tkacz, 1998). However, overall, research on such training effects remains scarce.

On the basis of correlational and training studies, to further clarify the mechanisms behind this transfer, neuroimaging evidence is needed. Previous research has revealed that both large- and small-scale spatial tasks recruit multiple cortical regions, such as the middle frontal gyrus, supplementary motor area (SMA), insula, and dorsolateral prefrontal cortex (dlPFC), which contribute to executive control, spatial integration, and decision-making (Patai & Spiers, 2021; Hawes et al., 2019). Both types of tasks also involve the superior parietal lobule (SPL) and inferior parietal lobule (IPL), which support the processing of spatial relationships between objects and/or the surrounding environment (Gogos et al., 2010; Calton & Taube, 2009). Large-scale spatial tasks uniquely activate the hippocampus for the organization of spatial memory (Eichenbaum, 2017), the parahippocampal gyrus for scene processing (Epstein, 2008; Baumann & Mattingley, 2021), and the posterior cingulate cortex for spatial information integration and updating (Clark et al., 2018; Wolbers et al., 2008). In contrast, small-scale spatial tasks more specifically engage the premotor cortex for motor planning (Doganci et al., 2023). Given that previous studies have typically investigated large- and small-scale spatial abilities separately, Li et al. (2019) conducted a meta-analysis of 103 fMRI studies and identified both overlapping and distinct activation patterns across the two scales. In line with previous findings, the commonly activated regions included the bilateral subgyrus, right superior frontal gyrus (SFG), right SPL, right superior/middle occipital gyrus (SOG/MOG), and bilateral precuneus. These regions are associated with visuospatial processing, working memory and executive control, potentially supporting shared cognitive processes across spatial scales. The analysis also revealed regions selectively activated by either large-scale tasks (e.g., the parahippocampal gyrus) or small-scale tasks (e.g., the inferior frontal gyrus, IFG), thus lending support to the view that the two abilities are partially dissociable.

Despite previous findings on the relationship between spatial abilities at different scales, important gaps remain. First, previous neuroimaging studies have focused primarily on either large- or small-scale spatial abilities, with few studies directly comparing both within the same participants. Second, evidence on cross-scale training transfer is still limited. Third, the neural mechanisms underlying potential training-related transfer across spatial scales remain unclear. In the present study, we conducted on-site navigation training in a campus environment, providing an ecologically valid context (Taube et al., 2013). Before and after training, participants performed two tasks each time: a distance judgment task assessing large-scale spatial ability and a mental rotation task assessing small-scale spatial ability. These tasks were selected because they are well-established representative tasks for each scale, and their stimuli and response can be matched across conditions to ensure comparability. Behavioral and neural differences between the two time points were compared across groups, which allowed the investigation of the training effect and its transfer. By combining real-world navigation training with brain imaging, this study moves beyond previous work that examined each spatial ability in isolation, offering new insights into how large- and small-scale spatial abilities are connected and how training can reshape the neural systems that support them.

## 2. Method

### 2.1. Participants

Thirty-two students from Beijing Normal University (13 males, 19 females, mean age 19.67 years, SD = 1.67) were randomly assigned to either a training group or a control group (16 participants per group, with no significant age or sex differences between the two groups). All participants were right-handed; had normal or corrected-to-normal vision; had no history of psychiatric, neurological, or brain injury conditions; and had lived on the campus for at least six months. All the participants signed an informed consent form before their participation and received appropriate compensation after the experiment. The study was approved by the Ethics Committee of Beijing Normal University. One participant from the training group was excluded because of excessive head motion (translation > 2 mm or rotation > 2°) during the small-scale task scans. This resulted in the use of brain imaging data from 31 participants (16 in the training group and 15 in the control group) for activation analysis. Additionally, six runs from five participants were excluded because of excessive head motion or failure to complete the task as required (no key responses for three or more consecutive trials). Brain activation for these participants was calculated on the basis of the remaining runs.

### 2.2. Experimental Procedure

Before training, all participants completed both large- and small-scale spatial ability tests while undergoing MRI scanning. Each scale’s task comprises four runs in a block design. The scanning sequence was as follows: two runs of the small-scale task, four runs of the large-scale task, and two runs of the small-scale task. This arrangement aimed to counterbalance task order and control for possible practice and fatigue effects. The day after the pretest, the training group began a continuous 20-day period of campus spatial navigation training. Within three days of completing the training or waiting period, both groups underwent the same posttest as the pretest, including MRI scanning of both tasks. The procedure is presented in Fig. 1.

**Fig. 1.**
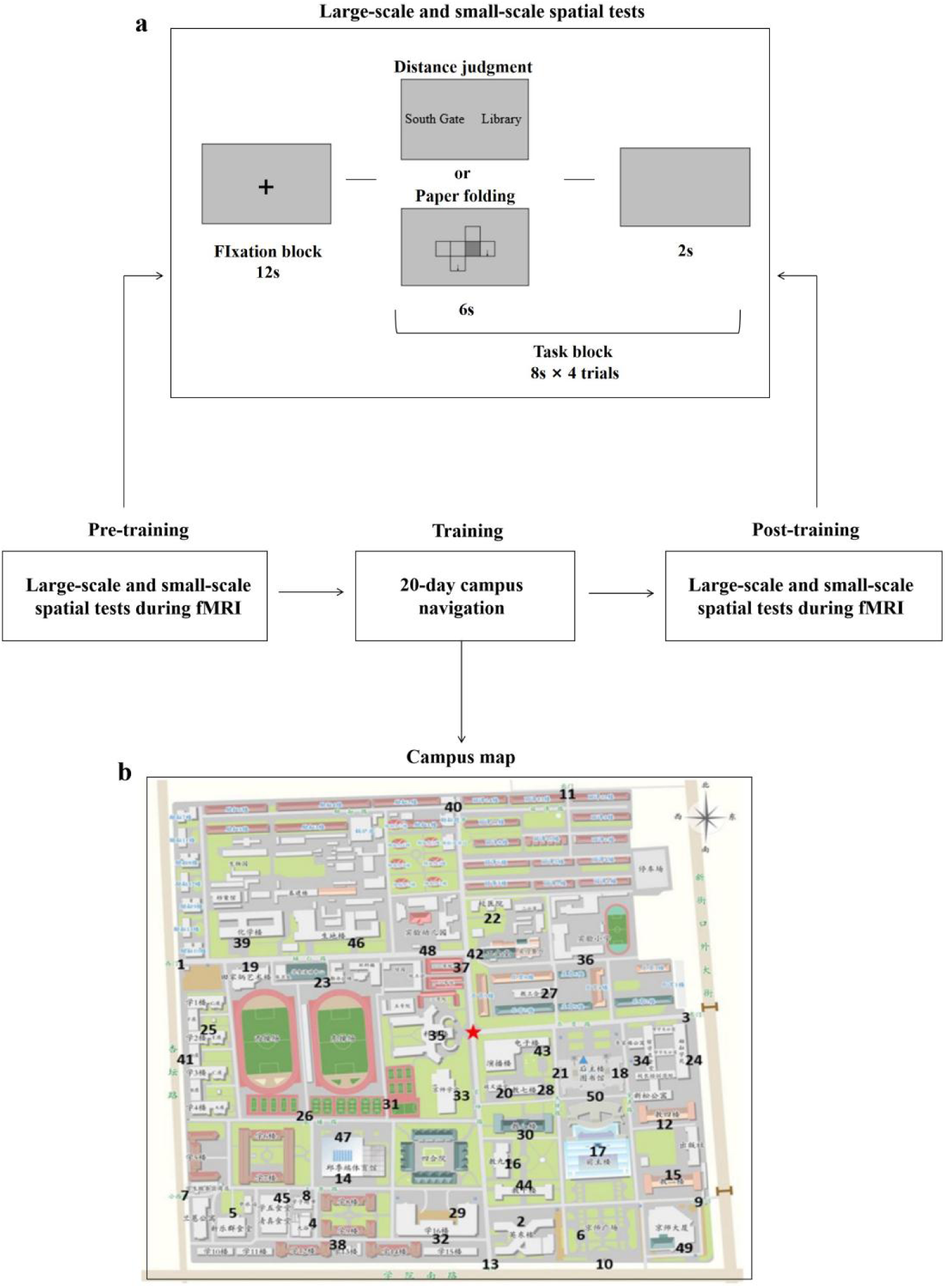
Experimental procedure. (a) Large-scale and small-scale spatial ability tests. Participants judged which of two locations (presented in Chinese characters in the real test) was closer to a target (distance judgment) or whether two edges would align when the diagram was folded into a cube (paper folding). This example shows one fixation block and one task block. (b) Campus map and target locations used for training and testing. The numbers indicate the 50 target locations, and the red star indicates the target location.

### 2.3. Large- and Small-Scale Spatial Ability Tests

Large-scale spatial ability was measured using a distance judgment task (Hirshhorn et al., 2012; Rosenbaum et al., 2004, 2005; Fig. 1a). The test began with a 12-second fixation block on the screen, after which two target campus locations were presented on the left and right sides of the screen for 6 seconds. Participants then had 8 seconds to judge which path from the two given locations to the predefined target location (marked by a star on the map, located centrally and familiar to participants) was shorter and to press the corresponding button. Failure to press the button within the time limit was considered an incorrect response. Each run consisted of seven fixation blocks and six task blocks presented alternately, with each task block including four distance judgments.

In the small-scale spatial ability test, the procedure and timing parameters matched those of the large-scale task, but the task was a paper folding task (Milivojevic et al., 2003; Shepard & Feng, 1972): a flat unfolded diagram of a cube was presented on the screen, with the dark square representing the cube’s base. Participants were asked to determine whether the two edges indicated by the arrows would match up when folded into a cube. They pressed the left key if they would align and the right key if they would not align.

### 2.4. Navigation Training

During the 20-day training period, participants completed 12-15 way-finding tasks daily at a minimum of three different locations. The training took approximately 30 minutes each day. In each training task, participants traveled to a specific location on campus and reported their GPS-based location to the experimenter via a messaging app. The experimenter then randomly selected one of 50 evenly distributed target locations (see Fig. 1b). Participants verbally described the shortest route from their location to the target location within one minute. The target locations given by the experimenter differed from those in the pre- and posttest tasks. After training, the participants’ route description accuracy increased to an average of 98.3%, with every participant achieving an accuracy rate above 90%.

### 2.5. MRI Scanning and Data Collection

MRI data were collected at the Brain Imaging Center of Beijing Normal University using a Siemens 3T MRI scanner (Siemens Trio) with a 12-channel head coil. Functional images were acquired using an echo-planar imaging (EPI) sequence with 33 slices (TR: 2,000 ms; TE: 30 ms; slices: 138; flip angle FA = 90°; voxel size: 3.125 × 3.125 × 3.5 mm³; FOV: 200 × 200 mm²). High-resolution structural T1-weighted images were simultaneously scanned using a 3D magnetization-prepared rapid gradient-echo (MP-RAGE) sequence (TR: 2530 ms; TE: 3.39 ms; TI: 1100 ms; slices: 128; flip angle FA = 7°; voxel size: 1 × 1 × 1.33 mm³). Earplugs were provided to reduce scanner noise interference, and participants were positioned supine on the scanner bed with their heads positioned in the center of the coil and secured with foam to minimize head movement. Each scan lasted 4 minutes and 36 seconds.

### 2.6. MRI Data Preprocessing and Statistical Analysis

The functional data were analyzed using the FEAT tool from FSL software (version 6.0; http://fsl.fmrib.ox.ac.uk/fsl/fslwiki/). Preprocessing steps included head motion correction, skull stripping, spatial smoothing (full width at half maximum of 5 mm), intensity normalization, high-pass filtering in the time domain (0.01 Hz), and registration of functional images to structural images and standard space.

For each run’s preprocessed data, statistical analysis of the time series was conducted using FMRIB’s Improved Linear Model (FILM). The onset times and durations of the stimuli were modeled and convolved with the hemodynamic response function (HRF). Six motion parameters (three translations and three rotations) were included as covariates of no interest in the first-level general linear model (GLM) to account for residual motion effects. Data from the four runs of each participant were combined using fixed-effects analyses, and the resulting data were entered into group-level mixed-effects analyses to account for both within- and between-subject variability. Task conditions (distance judgment and paper folding) were specified as explanatory variables (EVs) in the GLM. Task-related activation was estimated by contrasting each task EV against the baseline, producing activation maps for the two tasks.

To examine the effect of spatial navigation training on brain activation, we further used a mixed-effects model to compare pre- and posttest scans between the two groups. We considered only the changes in activation within the cortical gray matter (defined by the Harvard-Oxford template; Makris et al., 2006). Changes in brain activation were calculated as “(training group posttest - training group pretest) - (control group posttest - control group pretest)” to determine the training effect.

## 3. Results

### 3.1. Behavioral Results

To examine the relationship between large- and small-scale task performance, we conducted correlation analyses of reaction time and accuracy at the pretest. The results revealed a significant correlation for accuracy (r(29) = .40, p = .03) but not for reaction time (Fig. 2a). We next examined the effects of spatial navigation training using a mixed-design ANOVA with two factors: Group (training vs. control) and Time (pretest vs. posttest) (shown in Table 1 & Fig. 2b,c). For the large-scale task, we observed a significant main effect of time (F(1,30) = 16.50, p < .001, η_p_²=.36), a significant main effect of group (F(1,30) = 4.44, p = .04, η_p_²=.13) and an interaction effect of group×time (F(1,30) = 29.59, p < .001, η_p_² = .50) on accuracy. Post hoc tests revealed that the training group showed significantly greater accuracy in the posttest than in the pretest (F(1,30) = 45.14, p < .001, η_p_² = .60), whereas the control group showed no significant difference (F(1,30) = .95, p = .34). For the small-scale task, there was a significant main effect of time (F(1,30) = 20.12, p < .001, η_p_² = .40) but not of group (F(1,30)= .06, p = .81). The interaction effect between group and time was significant (F(1,30) = 9.01, p = .005, η_p_² = .23). Post hoc tests revealed that the training group had significantly greater accuracy in the posttest than in the pretest (F(1,30) = 28.02, p < .001, η_p_² = .48), whereas the control group did not significantly differ (F(1,30) = 1.10, p = .30). These results indicated that training enhanced performance accuracy in both large- and small-scale spatial tasks.

**Fig. 2.**
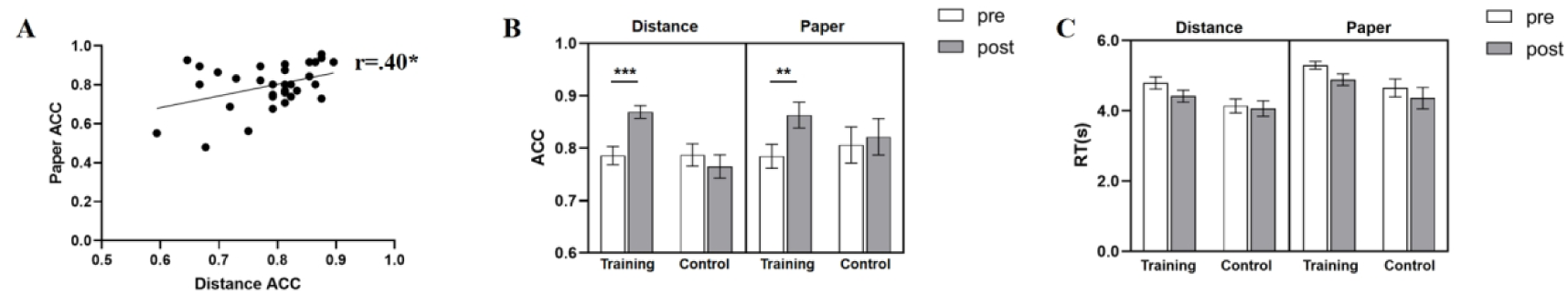
(a) Correlations between pretest performance in large-scale (distance judgment) and small-scale (paper folding) tasks. (b) Accuracy and (c) reaction times for both tasks across groups and sessions. The error bars represent ±1 standard error; *p < .05; **p < .01; ***p < .001.

**Table 1.** Accuracy and response time in the distance judgment and paper folding tasks (mean ± SD).

|  | Distance judgment |  |  |  | Paper folding |  |  |  |
| --- | --- | --- | --- | --- | --- | --- | --- | --- |
|  | ACC |  | RT(s) |  | ACC |  | RT(s) |  |
|  | pre | post | pre | post | pre | post | pre | post |
| Training group | 0.79<br>(0.07) | 0.87<br>(0.05) | 4.79<br>(0.55) | 4.42<br>(0.51) | 0.78<br>(0.09) | 0.86<br>(0.10) | 5.30<br>(0.45) | 4.89<br>(0.66) |
| Control group | 0.79<br>(0.09) | 0.76<br>(0.09) | 4.14<br>(0.61) | 4.07<br>(0.73) | 0.81<br>(0.14) | 0.82<br>(0.14) | 4.65<br>(1.01) | 4.36<br>(1.22) |

In addition to accuracy, we also examined response times to evaluate training effects. Although response times tended to improve after training, only the main effect of time in the small-scale task reached statistical significance in the small-scale task (F(1,30) = 30.00, p = .004). No other main effects or interactions reached significance.

### 3.2. Brain Activation in Large-Scale and Small-Scale Spatial Tasks

To compare brain activation patterns associated with different types of spatial abilities, we combined pretest brain imaging data from both groups—which were randomly assigned prior to training and thus were not expected to differ at baseline—and analyzed activations during the distance judgment and paper folding tasks. Statistical maps were thresholded at voxel-level p < .01 and cluster-level p < .05 (FDR-corrected) and visualized with BrainNet Viewer (Xia, Wang, & He, 2013; see Fig. 3). The results indicated that the large-scale and small-scale spatial tasks resulted in extensive common brain activation in regions including but not limited to the bilateral prefrontal cortex, posterior parietal cortex, and occipital visual areas. The hippocampus and parahippocampal region were activated only during large-scale tasks, whereas more widespread activation was detected in the temporoparietal and inferior frontal areas during small-scale tasks. The deactivation maps for both tasks are presented in Fig. S1 of the supplementary materials.

**Fig. 3.**
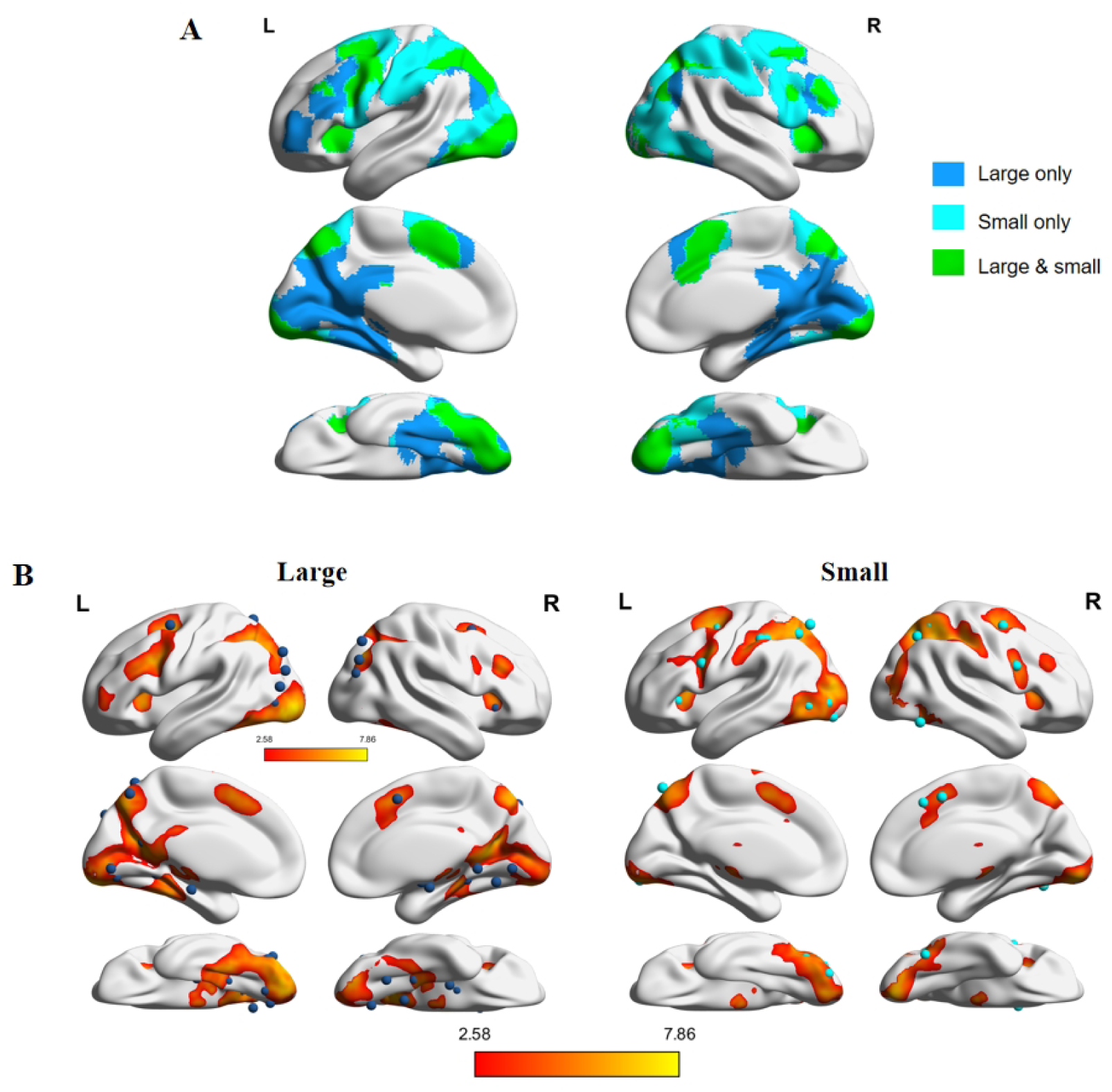
Brain activation in large- and small-scale spatial tasks at the pretest. (a) Significant activations thresholded at voxel-level p < .01 and cluster-level p < .05, corrected. Colors indicate regions activated exclusively in the large-scale task, exclusively in the small-scale task, or in both. (b) Significant activations in large-scale (left) and small-scale (right) tasks, with meta-analytic peaks from Li et al. (2019) shown as dark blue (large-scale) and light blue (small-scale) spheres.

To clearly distinguish individual activation clusters within each spatial task, a stringent statistical threshold (z > 5.20, p < 10^⁻7^) was applied. This threshold limited large contiguous activations, making it easier to identify distinct peak locations within broader regions. Regions that remained significantly activated under this threshold are shown in Table 2 and Table 3.

**Table 2.** Significant brain activation in large-scale spatial tasks at the pretest (p < 10^-7^). All coordinates are in MNI space; BA = Brodmann area; Z values indicate peak voxel statistics from the group-level analysis.

| Cluster | Brain Region | Hemisphere | BA | x | y | z | Max z-value | Voxels |
| --- | --- | --- | --- | --- | --- | --- | --- | --- |
| 1 | Middle Occipital Gyrus | L | 18 | -32 | -92 | -4 | 7.85 | 151 |
| 2 | Lingual Gyrus | R | 17 | 24 | -90 | -8 | 7.26 | 355 |
| 3 | Middle Frontal Gyrus | L | 6 | -30 | -2 | 50 | 7.05 | 158 |
| 4 | Precuneus | R | 7 | -28 | -70 | 40 | 6.97 | 399 |
| 5 | Posterior Cingulate | R | 19 | 18 | -60 | 18 | 6.80 | 398 |
| 6 | Superior Parietal Lobule | R | 7 | 12 | -72 | 52 | 6.76 | 146 |
| 7 | Middle Occipital Gyrus | R | 19 | 38 | -76 | 36 | 6.71 | 248 |
| 8 | Precuneus | L | 7 | -6 | -72 | 46 | 6.56 | 193 |
| 9 | Inferior Frontal Gyrus | L | 6 | -44 | 2 | 30 | 6.50 | 271 |
| 10 | Fusiform Gyrus | L | 37 | -36 | -46 | -18 | 6.43 | 1140 |
| 11 | Superior Occipital Gyrus | L | 19 | -32 | -78 | 26 | 6.36 | 567 |
| 12 | Supplementary Motor Area | L | 6 | 0 | 10 | 52 | 6.29 | 117 |
| 13 | Insula | L | 45 | -32 | 20 | 0 | 6.20 | 109 |
| 14 | Parahippocampal Gyrus | R | 36 | 24 | -32 | -14 | 5.72 | 22 |
| 15 | Middle Frontal Gyrus | R | 6 | 32 | 2 | 56 | 5.69 | 44 |

**Table 3.** Significant brain activation in small-scale spatial tasks at the pretest (p < 10^-7^). All coordinates are in MNI space; BA = Brodmann area; Z values indicate peak voxel statistics from the group-level analysis.

| Cluster | Brain Region | Hemisphere | BA | x | y | z | Max z-value | Voxels |
| --- | --- | --- | --- | --- | --- | --- | --- | --- |
| 1 | Superior Frontal Gyrus | L | 6 | -22 | -4 | 54 | 7.31 | 388 |
| 2 | Superior Parietal Lobule | L | 7 | -26 | -58 | 52 | 7.10 | 779 |
| 3 | Precuneus | L | 7 | -22 | -68 | 40 | 7.10 | 1684 |
| 4 | Inferior Parietal Lobule | R | 7 | 40 | -40 | 44 | 7.04 | 1849 |
| 5 | Lingual Gyrus | R | 18 | 12 | -90 | -10 | 6.68 | 296 |
| 6 | Lingual Gyrus | R | 17 | 12 | -90 | -10 | 6.68 | 33 |
| 7 | Inferior Frontal Gyrus | L | 6 | -46 | 2 | 28 | 6.60 | 160 |
| 8 | Middle Frontal Gyrus | R | 6 | 34 | 0 | 56 | 6.50 | 317 |
| 9 | Inferior Frontal Gyrus | R | 9 | 54 | 10 | 28 | 6.48 | 98 |
| 10 | Insula | L | 45 | -32 | 18 | 6 | 6.25 | 89 |
| 11 | Medial Frontal Gyrus | L | 6 | -4 | 6 | 52 | 5.75 | 47 |
| 12 | Insula | R | 45 | 32 | 20 | 6 | 5.51 | 29 |

### 3.3. Training Effects on Large-Scale Spatial Ability

To examine the effects of spatial navigation training on brain activation on the same scale, we compared pre- and posttest brain activation data for large-scale tasks between the two groups. We examined gray matter regions showing significant differences between the ‘training group posttest-training group pretest’ contrast and the ‘control group posttest-control group pretest’ contrast (voxel level: p < .01; cluster level: p < .05, corrected). This analysis revealed significant positive differences in activation in two brain regions (Fig. 4a), whereas no regions showed significant negative differences.

**Fig. 4.**
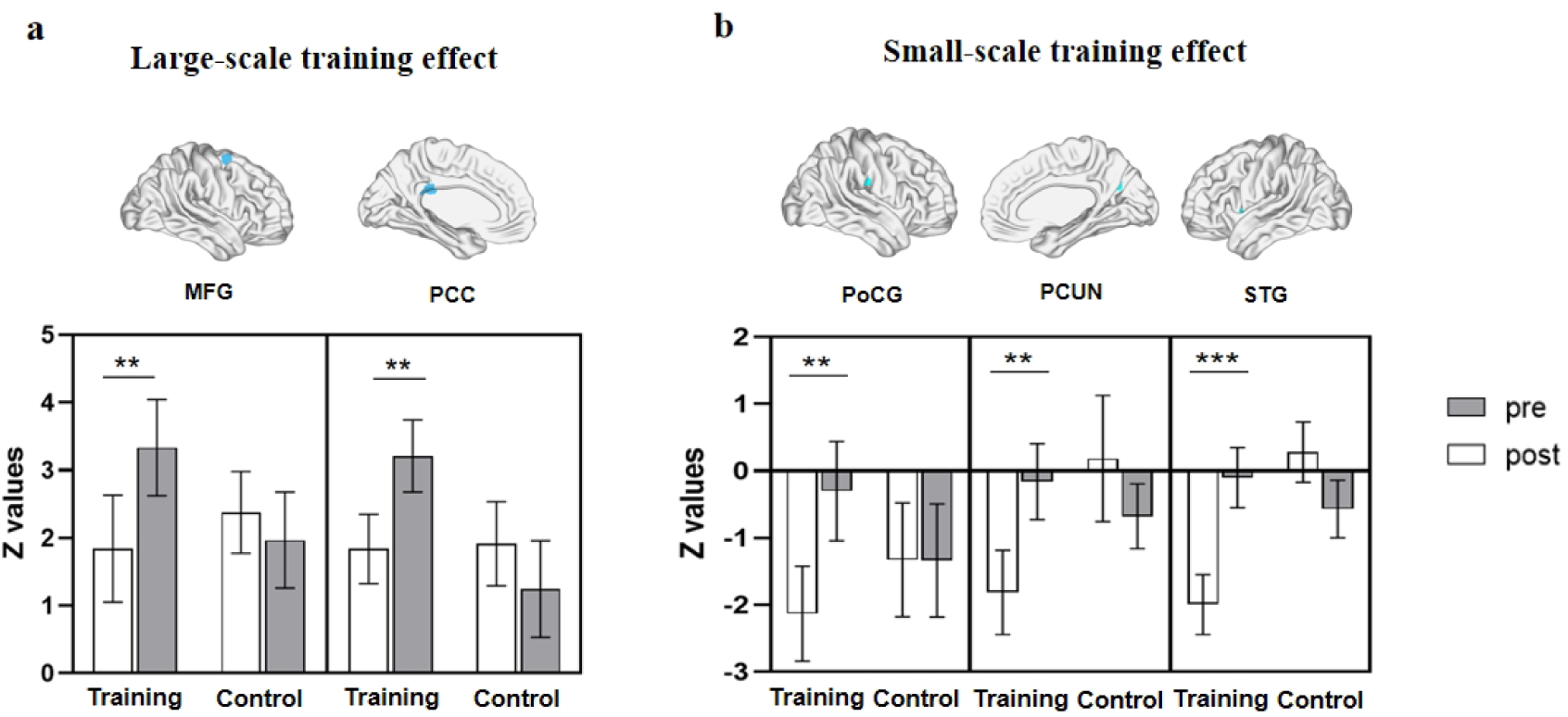
Brain regions showing significant differences in Z values between the training and control groups from pre- to posttraining during large-scale (a) and small-scale (b) spatial tasks, with corresponding signal values. MFG: middle frontal gyrus; PCC: posterior cingulate cortex; PoCG: postcentral gyrus; PCUN: precuneus; STG: superior temporal gyrus. The error bars represent ±1 standard error; **p < .01; ***p < .001.

One region was located in the right middle frontal gyrus and extended into the superior frontal gyrus (SFG) and subgyral region (48 voxels; peak coordinates: 26, 4, 66; Z_max_ = 3.51). The other one was in the bilateral posterior cingulate cortex (29 voxels; peak coordinates: −2, −32, 26; Z_max_ = 3.16). These regions overlapped with the activation observed during the distance judgment task at the pretest.

To better understand how navigation training affects brain activation within these significant regions, average signal changes were extracted from each area. In the training group, both regions exhibited a significant increase in activation from pretest to posttest (t_MFG_ = 3.08, p = .004, η_p_² = .23; t_PCC_ = 3.14, p = .004, η_p_² = .24), whereas no significant changes were observed in the control group (ps > .16). These findings indicate that navigation training enhanced positive activation in these regions during large-scale spatial tasks.

### 3.4. Training Effects on Small-Scale Spatial Ability

To investigate the transfer of large-scale training effects to brain activation during the small-scale task, we examined gray matter regions showing significant training effects for small-scale tasks (same contrast and threshold as large-scale tasks). We found significant negative differences in activation across three brain regions (Fig. 4b), with no significant positive differences observed.

One region was located in the right postcentral gyrus (26 voxels; peak coordinates: 58, −20, 28; Z_max_ = 3.40). Another one was in the right precuneus, extending into the subgyral region (28 voxels; peak coordinates: 16, −62, 28; Z_max_ = 3.38). The third one was in the left superior temporal gyrus (STG), extending into the insula (33 voxels; peak coordinates: −48, 6, 2; Z_max_ = 3.55). Among these regions, only the right precuneus overlapped with areas activated during the distance judgment task at the pretest; all three regions overlapped with pretest deactivation in the paper folding task.

With respect to average signal changes in the small-scale task, the three regions showed deactivation at the pretest. Following training, the degree of deactivation was reduced in the training group (t_PoCG_ = 2.87, p = .008, η_p_² = .21; t_PCUN_ = 3.49, p = .002, η_p_² = .28; t_STG_ = 4.72, p < .001, η_p_² = .42), whereas the control group showed no significant change (ps > .13). These results indicate that navigation training reduced deactivation in these three regions from pre- to posttraining during small-scale spatial tasks.

## 4. Discussion

This study examined the relationship between large- and small-scale spatial abilities by testing whether spatial navigation training is transferred to small-scale spatial performance. The results revealed a training effect, with the training group exhibiting greater improvements in accuracy than the control group did in both tasks. At the neural level, the training effect manifested as region-specific increases in activation during the large-scale task and decreases in deactivation during the small-scale task. The transfer of training effects indicates that the two spatial abilities are functionally related, whereas training-induced improvements in large- and small-scale abilities rely on distinct neural mechanisms.

Despite previous evidence that large- and small-scale spatial abilities are correlated, the cognitive mechanisms underlying this association remain unclear. We found that both spatial tasks elicited extensive overlapping brain activation, including in regions such as the bilateral dlPFC, PPC, and occipital visual areas, which is consistent with their role in visuospatial attention and processing (Pollmann & Yves von Cramon, 2000; Blankenburg et al., 2010). The hippocampus and parahippocampal gyrus were activated primarily during large-scale spatial tasks, which is consistent with previous findings on their role in spatial memory (Clark et al., 2018), a function that is mainly required in the distance judgment task. In contrast, small-scale tasks involve a broader network, including the temporoparietal and inferior frontal regions, associated with working memory and mental manipulation processes (Gogos et al., 2010; Kochan et al., 2011). These activation patterns are consistent with the meta-analysis by Li et al. (2019), indicating that large- and small-scale spatial abilities rely on both shared and distinct neural substrates.

To examine the training effect, we compared pre- and posttest task performance between groups. In the distance judgment task, the training group showed significant improvement in accuracy. Previous studies have reported that spatial navigation training induces changes in cortical thickness (Lövdén et al., 2012; Wenger et al., 2012) and in key navigation-related regions such as the hippocampus, parahippocampal cortex, and retrosplenial cortex (Woolley et al., 2015; Zhang et al., 2012; Auger et al., 2017). Our study extended this line of research by examining training effects across the whole brain and revealed training-related neural changes in the PCC and right MFG. These regions showed positive activation at the pretest, and this activation further increased after training. The MFG is associated with visual-spatial working memory (Desco et al., 2011), top-down mental manipulation (Hawes et al., 2019), and self-monitoring during imagery (De Lange et al., 2008). Previous studies also reported increased MFG activation after perceptual (Hötting et al., 2013) and alertness (Thimm et al., 2006) training during spatial tasks. We speculate that the navigation training in our study enhanced these general spatial cognitive components, together with spatial memory retrieval supported by the PCC (Ekstrom et al., 2011), which jointly improved distance judgment performance. This pattern indicates that large-scale spatial training may improve ability by increasing the involvement of perceptual and memory-related regions that extend beyond scene-processing areas.

Next, to assess cross-scale transfer, we compared paper folding task performance between the groups and found that the training group showed significantly higher posttest accuracy. This effect is unlikely to reflect general cognitive improvements; as such, improvements would be expected to produce similar activations across both tasks. Instead, training-related neural changes were specifically observed in the right postcentral gyrus, right precuneus, and left STG. These regions showed deactivation at the pretest, which was reduced after navigation training.

Previous studies reported that lower activation in these regions was associated with static or shape-based spatial processing, whereas they reported increased activation in dynamic or relational spatial tasks (Hurwitz et al., 2011; Manoach et al., 2004). We propose that the posttraining change in brain activation reflects a shift in strategy use during the small-scale spatial task. Burte et al. (2019) reported that participants may adopt several different strategies in paper folding tasks and that these strategies are linked to task performance. One possible explanation for our findings is that in the pretest paper folding task, participants mainly used a static and deliberate strategy. This is a strategy that relies on effortful analysis and may suppress mental imagery, leading to poorer performance. Navigation training provided participants with spatial experience, which encouraged a shift toward an imagery-based approach. This change engaged previously suppressed spatial processes and improved performance—for instance, by folding shapes in the mind to intuitively determine edge alignment. This hypothesis is further supported by a case study of a patient with no mental imagery (Zeman et al., 2010), who relied on a verbal strategy during spatial tasks and showed deactivation in the temporal gyrus. Nevertheless, this shift was not observed in the large-scale spatial task, as participants already favored visual strategies in such tasks (Markostamou et al., 2024).

Our findings show that spatial training affects a broader set of brain regions than those commonly activated for large- and small-scale tasks, with improvements in large-scale abilities and their transfer to small-scale tasks relying on distinct neural mechanisms. In the large-scale task, activation increased in the MFG and the PCC, both of which were already positively activated at the pretest. This increase reflects enhanced spatial cognition and memory functions following same-scale spatial training. In contrast, training-induced improvements in small-scale task performance, which indicate a relationship between both spatial abilities, were observed together with reduced inhibition in three brain regions. These notable results suggest that spatial abilities at different scales are integrated via mechanisms beyond shared spatial cognitive processes, most likely involving general problem-solving strategies and nonspatial components.

A key limitation of the present study lies in the absence of a nonspatial control task. Future work should include matched nonspatial imagery and working memory tasks to determine whether cross-scale transfer reflects neural changes specific to spatial tasks or general cognitive mechanisms. Other limitations include the following: First, although distance judgment and paper folding tests are commonly used to represent large- and small-scale spatial abilities, respectively, they differ not only in spatial scale but also in the type of knowledge and cognitive resources needed. This complexity makes ruling out confounding factors affecting the observed cross-scale training effects difficult. Second, the navigation training was conducted in a real-world environment, which, while ecologically valid, may have introduced variables such as first-person experience and environmental complexity, potentially affecting the training outcomes. Future studies should adopt experimental designs with greater experimental control—such as training conditions matched for physical activity demands but lacking navigational requirements—to minimize these confounding factors and isolate the specific contribution of spatial training.

## 5. Conclusion

The present study shows that large-scale and small-scale spatial cognition involve both shared and distinct neural activation. Large-scale spatial training is generalized to small-scale spatial tasks, further supporting the relationship between the two abilities. Nevertheless, small-scale spatial ability improvements manifested as reduced brain deactivation, in contrast to large-scale spatial ability, which showed increased positive activation after training. These similarities and differences support the partial dissociation model of large-scale and small-scale spatial abilities. They also provide a more comprehensive understanding of spatial cognition across scales and a conceptual basis for designing cross-scale interventions to enhance spatial skills.

## References

Auger, S. D., Zeidman, P., & Maguire, E. A. (2017). Efficacy of navigation may be influenced by retrosplenial cortex-mediated learning of landmark stability. Neuropsychologia, 104, 102–112. 10.1016/j.neuropsychologia.2017.08.012

Baumann, O., & Mattingley, J. B. (2021). Extrahippocampal contributions to spatial navigation in humans: A review of the neuroimaging evidence. Hippocampus, 31(7), 640–657. 10.1002/hipo.23313

Blankenburg, F., Ruff, C. C., Bestmann, S., Bjoertomt, O., Josephs, O., Deichmann, R., & Driver, J. (2010). Studying the role of human parietal cortex in visuospatial attention with concurrent TMS-fMRI. Cerebral Cortex, 20(11), 2702–2711. 10.1093/cercor/bhq015

Burte, H., Gardony, A. L., Hutton, A., & Taylor, H. A. (2019). Knowing when to fold’em: Problem attributes and strategy differences in the Paper Folding test. Personality and Individual Differences, 146, 171–181. 10.1016/j.paid.2018.08.009

Calton, J. L., & Taube, J. S. (2009). Where am I and how will I get there from here? A role for posterior parietal cortex in the integration of spatial information and route planning. Neurobiology of Learning and Memory, 91(2), 186–196. 10.1016/j.nlm.2008.09.015

Clark, B. J., Simmons, C. M., Berkowitz, L. E., & Wilber, A. A. (2018). The retrosplenial-parietal network and reference frame coordination for spatial navigation. Behavioral Neuroscience, 132(5), 416–429. 10.1037/bne0000260

De Lange, F. P., Roelofs, K., & Toni, I. (2008). Motor imagery: a window into the mechanisms and alterations of the motor system. Cortex, 44(5), 494–506. 10.1016/j.cortex.2007.09.002

Desco, M., Navas-Sánchez, F. J., Sánchez-González, J., Reig, S., Robles, O., Franco, C., Guzmán-De-Villoria, J. A., García-Barreno, P., & Arango, C. (2011). Mathematically gifted adolescents use more extensive and more bilateral areas of the fronto-parietal network than controls during executive functioning and fluid reasoning tasks. NeuroImage, 57(1), 281–292. 10.1016/j.neuroimage.2011.03.063

Doganci, N., Iannotti, G. R., Coll, S. Y., & Ptak, R. (2023). How embodied is cognition? fMRI and behavioral evidence for common neural resources underlying motor planning and mental rotation of bodily stimuli. Cerebral Cortex, 33(22), 11146–11156. 10.1093/cercor/bhad352

Eichenbaum, H. (2017). The role of the hippocampus in navigation is memory. Journal of Neurophysiology, 117(4), 1785–1796. 10.1152/jn.00005.2017

Ekstrom, A. D., Copara, M. S., Isham, E. A., Wang, W. C., & Yonelinas, A. P. (2011). Dissociable networks involved in spatial and temporal order source retrieval. NeuroImage, 56(3), 1803–1813. 10.1016/j.neuroimage.2011.02.033

Epstein, R. A. (2008). Parahippocampal and retrosplenial contributions to human spatial navigation. Trends in Cognitive Sciences, 12(10), 388–396. 10.1016/j.tics.2008.07.004

Evans, G. W. (1980). Environmental cognition. Psychological Bulletin, 88(2), 259–287. 10.1037/0033-2909.88.2.259

Farr, A. C., Kleinschmidt, T., Yarlagadda, P., & Mengersen, K. (2012). Wayfinding: A simple concept, a complex process. Transport Reviews, 32(6), 715–743. 10.1080/01441647.2012.712555

Gogos, A., Gavrilescu, M., Davison, S., Searle, K., Adams, J., Rossell, S. L., Bell, R., Davis, S. R., & Egan, G. F. (2010). Greater superior than inferior parietal lobule activation with increasing rotation angle during mental rotation: An fMRI study. Neuropsychologia, 48(2), 529–535. 10.1016/j.neuropsychologia.2009.10.013

Hawes, Z., Sokolowski, H. M., Ononye, C. B., & Ansari, D. (2019). Neural underpinnings of numerical and spatial cognition: An fMRI meta-analysis of brain regions associated with symbolic number, arithmetic, and mental rotation. Neuroscience & Biobehavioral Reviews, 103, 316–336. 10.1016/j.neubiorev.2019.05.007

Hegarty, M., Montello, D. R., Richardson, A. E., Ishikawa, T., & Lovelace, K. (2006). Spatial abilities at different scales: Individual differences in aptitude-test performance and spatial-layout learning. Intelligence, 34(2), 151–176. 10.1016/j.intell.2005.09.005

Hirshhorn, M., Grady, C., Rosenbaum, R. S., Winocur, G., & Moscovitch, M. (2012). The hippocampus is involved in mental navigation for a recently learned, but not a highly familiar environment: A longitudinal fMRI study. Hippocampus, 22(4), 842–852. 10.1002/hipo.20944

Hötting, K., Holzschneider, K., Stenzel, A., Wolbers, T., & Röder, B. (2013). Effects of a cognitive training on spatial learning and associated functional brain activations. BMC Neuroscience, 14(1), 73. 10.1186/1471-2202-14-73

Hurwitz, M., Valadao, D., & Danckert, J. (2011). Functional MRI of dynamic judgments of spatial extent. Experimental Brain Research, 214(1), 61–72. 10.1007/s00221-011-2806-9

Jansen, P., Wiedenbauer, G., & Hahn, N. (2010). Manual rotation training improves direction-estimations in a virtual environmental space. European Journal of Cognitive Psychology, 22(1), 6–17. 10.1080/09541440802678487

Kochan, N. A., Valenzuela, M., Slavin, M. J., McCraw, S., Sachdev, P. S., & Breakspear, M. (2011). Impact of load-related neural processes on feature binding in visuospatial working memory. PLoS One, 6(8), e23960. 10.1371/journal.pone.0023960

Kozhevnikov, M., & Hegarty, M. (2001). A dissociation between object manipulation spatial ability and spatial orientation ability. Memory & Cognition, 29, 745–756. 10.3758/BF03200477

Li, Y., Kong, F., Ji, M., Luo, Y., Lan, J., & You, X. (2019). Shared and distinct neural bases of large-and small-scale spatial ability: a coordinate-based activation likelihood estimation meta-analysis. Frontiers in Neuroscience, 12, 1021. 10.3389/fnins.2018.01021

Likhanov, M., Maslennikova, E., Costantini, G., Budakova, A., Esipenko, E., Ismatullina, V., & Kovas, Y. (2022). This is the way: Network perspective on targets for spatial ability development programmes. British Journal of Educational Psychology, 92(4), 1597–1620. 10.1111/bjep.12524

Lövdén, M., Schaefer, S., Noack, H., Bodammer, N. C., Kühn, S., Heinze, H.-J., Lindenberger, U. (2012). Spatial navigation training protects the hippocampus against age-related changes during early and late adulthood. Neurobiology of Aging, 33(3), 620.e9–620.e22. 10.1016/j.neurobiolaging.2011.02.013

Makris, N., Goldstein, J. M., Kennedy, D., Hodge, S. M., Caviness, V. S., Faraone, S. V., Tsuang, M. T., & Seidman, L. J. (2006). Decreased volume of left and total anterior insular lobule in schizophrenia. Schizophrenia Research, 83(2-3), 155–171. 10.1016/j.schres.2005.11.020

Manoach, D. S., White, N. S., Lindgren, K. A., Heckers, S., Coleman, M. J., Dubal, S., & Holzman, P. S. (2004). Hemispheric specialization of the lateral prefrontal cortex for strategic processing during spatial and shape working memory. NeuroImage, 21(3), 894–903. 10.1016/j.neuroimage.2003.10.025

Markostamou, I., Morrissey, S., & Hornberger, M. (2024). Imagery and verbal strategies in spatial memory for route and survey descriptions. Brain Sciences, 14(4), 403. 10.3390/brainsci14040403

Milivojevic, B., Johnson, B. W., Hamm, J. P., & Corballis, M. C. (2003). Non-identical neural mechanisms for two types of mental transformation: Event-related potentials during mental rotation and mental paper folding. Neuropsychologia, 41(10), 1345–1356. 10.1016/S0028-3932(03)00060-5

Patai, E. Z., & Spiers, H. J. (2021). The versatile wayfinder: prefrontal contributions to spatial navigation. Trends in Cognitive Sciences, 25(6), 520–533. 10.1016/j.tics.2021.02.010

Pollmann, S., & Yves von Cramon, D. (2000). Object working memory and visuospatial processing: functional neuroanatomy analyzed by event-related fMRI. Experimental Brain Research, 133(1), 12–22. 10.1007/s002210000396

Quaiser-Pohl, C., Lehmann, W., & Eid, M. (2004). The relationship between spatial abilities and representations of large-scale space in children—a structural equation modeling analysis. Personality and Individual Differences, 36(1), 95–107. 10.1016/S0191-8869(03)00071-0

Rosenbaum, R. S., Gao, F., Richards, B., Black, S. E., & Moscovitch, M. (2005). “Where to?” remote memory for spatial relations and landmark identity in former taxi drivers with Alzheimer’s disease and encephalitis. Journal of Cognitive Neuroscience, 17(3), 446–462. 10.1162/0898929053279496

Rosenbaum, R. S., Ziegler, M., Winocur, G., Grady, C. L., & Moscovitch, M. (2004). “I have often walked down this street before”: fMRI studies on the hippocampus and other structures during mental navigation of an old environment. Hippocampus, 14(7), 826–835. 10.1002/hipo.10218

Shepard, R. N., & Feng, C. (1972). A chronometric study of mental paper folding. Cognitive Psychology, 3(2), 228–243. 10.1016/0010-0285(72)90005-9

Taube, J. S., Valerio, S., & Yoder, R. M. (2013). Is navigation in virtual reality with FMRI really navigation?. Journal of Cognitive Neuroscience, 25(7), 1008–1019. 10.1162/jocn_a_00386

Thimm, M., Fink, G. R., Küst, J., Karbe, H., & Sturm, W. (2006). Impact of alertness training on spatial neglect: a behavioural and fMRI study. Neuropsychologia, 44(7), 1230–1246. 10.1016/j.neuropsychologia.2005.09.008

Tkacz, S. (1998). Learning map interpretation: Skill acquisition and underlying abilities. Journal of Environmental Psychology, 18(3), 237–249. 10.1006/jevp.1998.0094

Uttal, D. H., Meadow, N. G., Tipton, E., Hand, L. L., Alden, A. R., Warren, C., & Newcombe, N. S. (2013). The malleability of spatial skills: a meta-analysis of training studies. Psychological Bulletin, 139(2), 352. 10.1037/a0028446

Walter, E., & Dassonville, P. (2011). Activation in a frontoparietal cortical network underlies individual differences in the performance of an embedded figures task. PloS One, 6(7), e20742. 10.1371/journal.pone.0020742

Wang, L., Cohen, A. S., & Carr, M. (2014). Spatial ability at two scales of representation: A meta-analysis. Learning and Individual Differences, 36, 140–144. 10.1016/j.lindif.2014.10.006

Wenger, E., Schäfer, S., Noack, H., Kühn, S., Mårtensson, J., Heinze, H.-J., Düzel, E., Bäckmann, L., Lindenberger, U., & Lövden, M. (2012). Cortical thickness changes following spatial navigation training in adulthood and aging. NeuroImage, 59(4), 3389–3397. 10.1016/j.neuroimage.2011.11.015

Wexler, M., Kosslyn, S. M., & Berthoz, A. (1998). Motor processes in mental rotation. Cognition, 68(1), 77–94. 10.1016/S0010-0277(98)00032-8

Wolbers, T., Hegarty, M., Büchel, C., & Loomis, J. M. (2008). Spatial updating: how the brain keeps track of changing object locations during observer motion. Nature Neuroscience, 11(10), 1223–1230. 10.1038/nn.2189

Wolbers, T., & Wiener, J. M. (2014). Challenges for identifying the neural mechanisms that support spatial navigation: the impact of spatial scale. Frontiers in Human Neuroscience, 8, 571. 10.3389/fnhum.2014.00571

Woolley, D. G., Mantini, D., Coxon, J. P., D’Hooge, R., Swinnen, S. P., & Wenderoth, N. (2015). Virtual water maze learning in human increases functional connectivity between posterior hippocampus and dorsal caudate. Human Brain Mapping, 36(4), 1265–1277. 10.1002/hbm.22700

Xia, M., Wang, J., & He, Y. (2013). BrainNet Viewer: a network visualization tool for human brain connectomics. PloS One, 8(7), e68910.

Zeman, A. Z., Della Sala, S., Torrens, L. A., Gountouna, V. E., McGonigle, D. J., & Logie, R. H. (2010). Loss of imagery phenomenology with intact visuo-spatial task performance: A case of ‘blind imagination’. Neuropsychologia, 48(1), 145–155. 10.1016/j.neuropsychologia.2009.08.024

Zhang, H., Copara, M., & Ekstrom, A. D. (2012). Differential recruitment of brain networks following route and cartographic map learning of spatial environments. PLoS One, 7(9), e44886. 10.1371/journal.pone.0044886

