## Supplementary Material for "Spatial Navigation Training Enhances Large-Scale and Small-Scale Spatial Abilities through Different Neural Mechanisms"

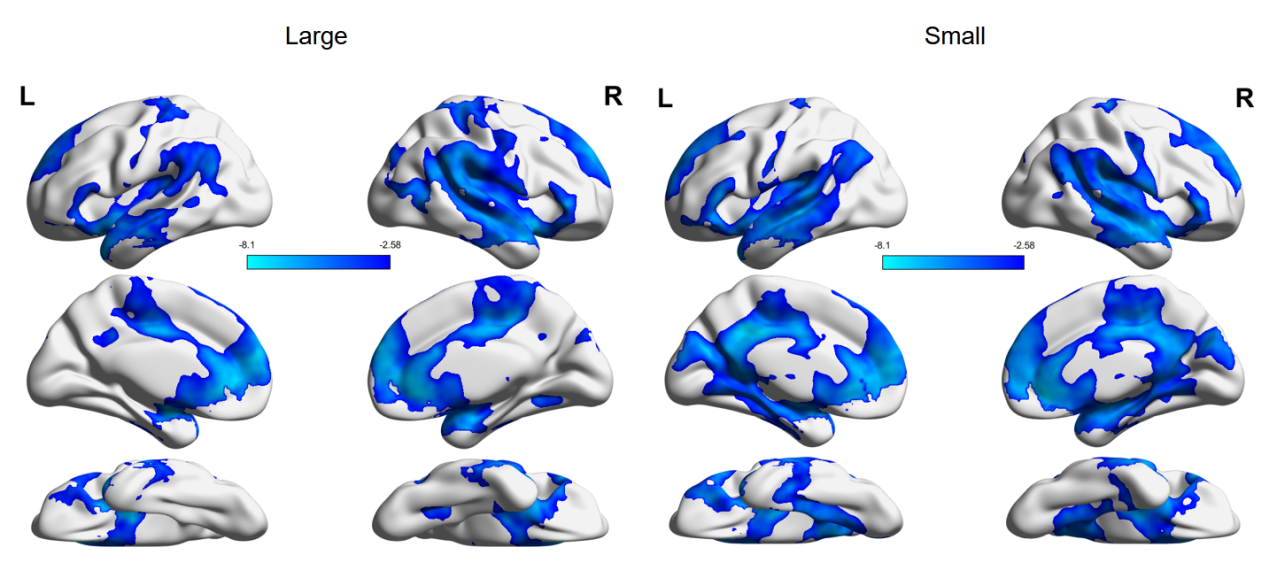

****Fig. S1.**** Brain deactivation in large- and small-scale spatial tasks at the pre-test, thresholded at a voxel-level of p < .01.
